# A combined computational and experimental investigation of the filtration function of splenic macrophages in sickle cell disease

**DOI:** 10.1101/2023.05.31.543007

**Authors:** Guansheng Li, Yuhao Qiang, He Li, Xuejin Li, Pierre A. Buffet, Ming Dao, George Em Karniadakis

## Abstract

Being the largest lymphatic organ in the body, the spleen also constantly controls the quality of red blood cells (RBCs) in circulation through its two major filtration components, namely interendothelial slits (IES) and red pulp macrophages. In contrast to the extensive studies in understanding the filtration function of IES, there are relatively fewer works on investigating how the splenic macrophages retain the aged and diseased RBCs, i.e., RBCs in sickle cell disease (SCD). Herein, we perform a computational study informed by companion experiments to quantify the dynamics of RBCs captured and retained by the macrophages. We first calibrate the parameters in the computational model based on microfluidic experimental measurements for sickle RBCs under normoxia and hypoxia, as those parameters are not available in the literature. Next, we quantify the impact of a set of key factors that are expected to dictate the RBC retention by the macrophages in the spleen, namely, blood flow conditions, RBC aggregation, hematocrit, RBC morphology, and oxygen levels. Our simulation results show that hypoxic conditions could enhance the adhesion between the sickle RBCs and macrophages. This, in turn, increases the retention of RBCs by as much as five-fold, which could be a possible cause of RBC congestion in the spleen of patients with SCD. Our study on the impact of RBC aggregation illustrates a ‘clustering effect’, where multiple RBCs in one aggregate can make contact and adhere to the macrophages, leading to a higher retention rate than that resulting from RBC-macrophage pair interactions. Our simulations of sickle RBCs flowing past macrophages for a range of blood flow velocities indicate that the increased blood velocity could quickly attenuate the function of the red pulp macrophages on detaining aged or diseased RBCs, thereby providing a possible rationale for the slow blood flow in the open circulation of the spleen. Furthermore, we quantify the impact of RBC morphology on their tendency to be retained by the macrophages. We find that the sickle and granular-shaped RBCs are more likely to be filtered by macrophages in the spleen. This finding is consistent with the observation of low percentages of these two forms of sickle RBCs in the blood smear of SCD patients. Taken together, our experimental and simulation results aid in our quantitative understanding of the function of splenic macrophages in retaining the diseased RBCs and provide an opportunity to combine such knowledge with the current knowledge of the interaction between IES and traversing RBCs to apprehend the complete filtration function of the spleen in SCD.

## 1. Introduction

The spleen, a dark red to blue-black elongated organ, is located in the abdomen and adjacent to the greater curvature of the stomach and within the omentum [1]. As the most significant secondary immune organ in the human body, the spleen prevents pathogenic microorganisms from remaining and multiplying in the bloodstream by initiating immune reactions [2–4]. Another essential role of the spleen is to clear red blood cells (RBCs) from circulation when their biochemical or biomechanical properties hinder their circulation functionality. The two physiological functions of the spleen are accomplished within its two functionally and morphologically distinct compartments, namely white pulp and red pulp [2, 4, 5]. The white pulp comprises three subcompartments: the periarteriolar lymphoid sheath (PALS), the follicles, and the marginal zone (MZ). While the PALS contains central arterioles surrounded predominately by the T cells, the follicles and the MZ are occupied mainly by B cells [3, 6–8]. After passing through the white pulp, blood flows into the medium-sized central artery from the splenic artery and then engages in either fast or slow microcirculation. As much as 90% of the splenic blood travels from the perifollicular zone to the venous sinus lumen through the fast microcirculation pathway [2, 9], whereas only 10% of the blood flows through the slow microcirculation, where the RBCs need to traverse the red pulp, a three-dimensional meshwork of splenic cords and venous sinuses, which works like a blood filter that removes damaged and aged RBCs [3, 10, 11]. Before returning to the vascular beds, RBCs have to squeeze through the narrow slits between endothelial cells (IES) in the wall of sinuses, the most stringent challenge for the deformability of RBCs in microcirculation, resulting in the retention of less deformable RBCs or the removal of intraerythrocytic bodies [12–14].

In the red pulp, RBCs also make close contact with abundant splenic macrophages, which are capable of detecting the deleterious changes in the RBC membrane and engulfing the aged or damaged RBCs through erythrophagocytosis [3, 15, 16]. The key factors that determine the potential targets of phagocytosis have been considered to be the biochemical signaling triggered by the interaction between the ligands on the RBCs and the receptors on the macrophages, such as binding of antibodies (NAbs) to the band-3 proteins [17, 18], increased exposure of PS [19, 20], decreased expression of CD47 [21, 22] and conformational changes in CD47 [21, 23]. It is thought that this ligand-receptor interaction could generate so-called “eat me” signals and provoke RBC clearance by macrophages in the spleen. Recently, the impact of biophysical properties of the targeted cells and the macrophages are also investigated. An *in vitro* experimental investigation [24] suggested that the impact of RBC rigidity could override that of CD47 in the process of phagocytosis, indicating that less deformable RBCs are more likely to be engulfed by splenic macrophages. This finding was echoed by a subsequent study [25] showing a high propensity of macrophages to recognize and phagocytose the lysed and less deformable RBCs over intact ones.

Although the function of the spleen in RBC physiology has been well studied, its role in RBC disorders is not yet fully understood. For example, in sickle cell disease (SCD), the mutant sickle hemoglobin (HbS) could polymerize into stiff fiber bundles under hypoxia, causing the RBCs to sickle [26, 27]. Sickling of RBCs induces drastic alterations of the RBC rigidity and leads to their distorted heterogeneous shapes, which are categorized into elongated, granular, oval, holly-leaf, and crescent (classic sickle) shapes [28–30]. In addition to their increased stiffness, sickle RBCs are featured with enhanced cell adhesion, contributing to the initiation and propagation of vaso-occlusion events, a hallmark of SCD [31–33]. The spleen is one of the most common and early organs to be affected in SCD. In SCD infants with still functional spleen, these sickle RBCs are amenable to mechanical retention by the spleen, a process that likely contributes to hemolytic anemia [34] and triggers acute splenic sequestration crises (ASSC), a life-threatening complication of SCD [35–41]. Children with SCD between five months and two years of age have a higher risk for ASSC, manifested as an abrupt fall in the hemoglobin level and splenomegaly [38, 42–46]. SCD patients with multiple episodes of ASSC require surgical splenectomy, a supportive transfusion program, or both. They are at higher risk of complications [47–49]. To date, the mechanisms causing ASSC remain elusive. An improved understanding of the filtration function of the spleen in SCD could advance our knowledge of this early complication of SCD.

Significant efforts have been accumulated to understand the filtration function of splenic IES in blood diseases. Several ex vivo [50–54] experiments were conducted through perfusing malaria-infected RBCs and biochemically treated RBCs that resemble the diseased RBCs through the donated human spleen and reported that RBCs with decreased deformability could be mechanically retained at IES. These findings were confirmed by a number of *in vitro* investigations [54–59], [ADD REF: S. Huang, et al., Infection and Immunity, 82 (2014) 2532-2541], where microfluidic chips were devised to mimic the filtration function of IES in blood diseases such as SCD, malaria, and spherocytosis. On the other hand, numerous computational studies have been performed to simulate the dynamics of RBCs passing through the IES and quantify the critical factors that dictate the passage of the RBCs, such as the surface-to-volume ratio of RBCs, the shear and bending modulus of the cells, RBC membrane viscosity, and the size of the IES[4, 60–66]. These simulations have complimented the *ex vivo* and *in vitro* experiments by providing the analysis of the deformation of RBCs, the shear strain and stress on the RBC membrane as well as the potential vesiculation and lysis of RBCs during their traversal through IES, which could not be directly observed from the experiments.

In contrast to the extensive literature on exploring the function of IES in clearing the aged and diseased RBCs in the spleen, there is a lack of studies on quantifying the filtration function of the splenic macrophages. A quantitative understanding of the role of macrophages in removing the RBCs will open the way to integrate the two major filtration components in the spleen that constantly control the quality of circulating RBCs and advance our knowledge of the underlying mechanism causing the splenic complications in SCD. Thus, in this work, we perform an integrated experimental and computational study to explore the clearance of RBC suspension by macrophages under physiologically relevant flow conditions in the spleen. We will quantify the influence of the velocity, hematocrit, and various shapes of sickle RBCs on the clearance efficiency under splenic conditions, which will be combined with our current understanding of the filtration function of IES to postulate the complete filtration function of the spleen in SCD.

## 2. Methods and materials

### 2.1. Computational methods and models

#### 2.1.1. DPD method and cellular level blood cell models

Several computational models of RBCs have been developed in the last two decades to simulate the dynamics of normal and diseased RBCs. Based on their level of complexity, these RBC models can be categorized into the protein-level RBC models [67–77], which are widely used in simulating the pathological alterations of RBC membrane structure in blood disorders, and cellular-level RBC models [72, 78–85], which are mostly used in modeling blood cell suspensions or blood flow. Due to the high computational cost of the protein-level RBC models, we employ a cellular-level model [86] developed based on dissipative particle dynamics (DPD) [87] to simulate the normal and sickle RBCs as well as macrophages (more details on the cellular-level models and the values of model parameters can be found in S1 Text). The DPD method is a mesoscopic particle-based simulation technique, where each DPD particle represents a lump of molecules and interacts with other particles through soft pairwise forces [88, 89]. DPD can provide the correct hydrodynamic behavior of fluids at the mesoscale, and it has been successfully applied to study complex fluids [90–95]. The formulation of the DPD method can be found in S1 Text.

#### 2.1.2. Stochastic adhesion model

To characterize the initial adhesion between RBCs and macrophages, we employ the stochastic bond formation/dissociation model proposed by Hammer and Apte [96, 97], where the adhesive bond can be formed and dissociated with predefined possibilities *P_on_*and *P_off_* within a critical length *d_on_*and *d_off_*, respectively [96] and they can be calculated by

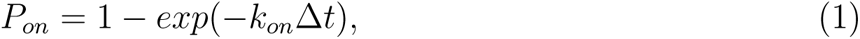

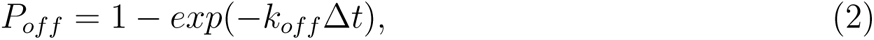

where Δ t is the timestep of the simulation; *k_on_* and *k_off_* are the formation and dissociation rates defined as:

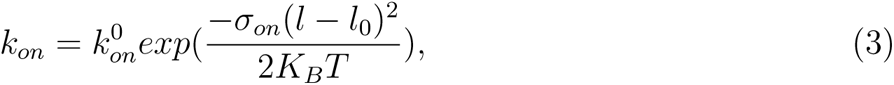

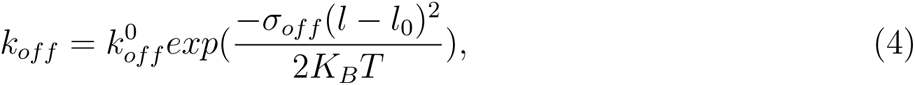

where *T* is the temperature, and 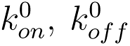 are the formation and dissociation rates at the equilibrium distance *l*_0_. *σ_on_* and *σ_off_* are the effective formation and rupture strengths within a given reactive distance *d_on_* and *d_off_*, respectively. For a given bond, an adhesion potential energy is given by 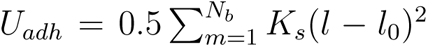, and the adhesive force between cells and macrophages is calculated by 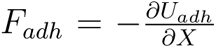, where *K_s_*is the adhesion strength, *N_b_* is the number of the formed bonds, *l* is the bond length. The following two conditions are used to determine the formation of a bond

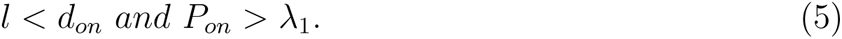

On the other hand, the following two conditions lead to the dissociation of a bond,

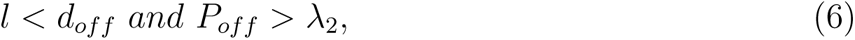

where the *λ*_1_ and *λ*_2_ are random numbers drawn from a uniform distribution between [0,1]. The values of the parameters used in the adhesion models and the associated references are summarised in Table 1.

**Table 1:** Model parameters for adhesion dynamics between RBCs and macrophages.

| Parameters | Simulation | Physical | Reported Range |
| --- | --- | --- | --- |
| $K_s$ | 50~150 | $(2.4 \sim 7.2)\mu N/m$ | $(1 \sim 10^4)\mu N/m$ [96, 98–100] |
| $l_0$ | 0.3 | $0.3\mu m$ | $(0.0 \sim 0.2)\mu m$ [92, 101, 102] |
| $d_{on}$ | 0.75 | $0.75\mu m$ | $(0.0 \sim 1.0)\mu m$ [92, 99–101] |
| $d_{off}$ | 0.75 | $0.75\mu m$ | $(0.0 \sim 1.0)\mu m$ [92, 99, 100, 103] |
| $\sigma_{on}$ | 10.58 | $0.51\mu N/m$ | $(0.5 \sim 0.6)\mu N/m$ [101] |
| $\sigma_{off}$ | 3.49 | $0.17\mu N/m$ | $(0.05 \sim 0.75)\mu N/m$ [101] |
| $k_{on}$ | 100 ~ 150 | $(5.42 \sim 8.14) * 10^4 s^{-1}$ | $(0.001 \sim 1.2) * 10^6 s^{-1}$ [100, 104] |
| $k_{off}$ | 0.001 ~ 0.01 | $0.54 \sim 5.42 s^{-1}$ | $(0.5 \sim 300) s^{-1}$ [105] |

### 2.2. Experimental and Simulation Setups

#### 2.2.1. Experimental setup

Fig. 1(a) illustrates the experimental setup and the observation of cell-macrophage adhesion. Healthy blood samples were acquired from the local blood bank, while sickle blood samples were obtained from homozygous (HbSS) sickle cell disease (SCD) patients at Massachusetts General Hospital. The collection of sickle blood samples followed the guidelines of an Excess Human Material Protocol, which was approved by the Partners Healthcare Institutional Review Board (IRB) with a waiver of consent. Additional HbSS blood samples were drawn from SCD patients at the University of Pittsburgh, following the University of Pittsburgh IRB protocol PRO08110422. In vitro microfluidic experiments were conducted under an approved exempt protocol (Massachusetts Institute of Technology IRB protocol E-1523). All experiments were conducted within 5 days of blood collection. Before the microfluidics experiments, the red blood cells were gently washed twice with phosphate buffered saline (PBS) solution at 2,000 rpm for 2 min at room temperature (20°C). The hematocrit of the RBC suspensions used within the microfluidic channels is estimated to be 1.0–2%. Before the adhesion experiments, RBCs were resuspended in PBS (2%, v/v), and were opsonized by incubation with 0.5*µM* IgG (Rockland, Limerick, PA, USA) at room temperature (20°C). THP-1 macrophages (ATCC, Manassas, VA, USA) were cultured in RPMI media (Stem-Cell Technologies, Cambridge, MA, USA) supplemented with 10% FBS (Sigma-Aldrich, St. Louis, MO, USA), and were differentiated for 2 days using 100 ng/ml phorbol myristate acetate (PMA) (StemCell Technologies, Cambridge, MA, USA). To study the adhesion dynamics of sickle RBCs under shear flow and hypoxia, we conducted experiments using a specially developed hypoxic microfluidic device [106]. This device consists of a dual-layer microchannel construction with a gas-permeable polydimethylsiloxane (PDMS) membrane (150 mm thick) and and two microchannels: a ‘flow microchannel’ for the RBC suspension and a ‘gas microchannel’ for delivering the desired gas mixture of carbon dioxide (*CO*_2_), oxygen (*O*_2_), and nitrogen (*N*_2_). Deoxygenation and oxygenation conditions of the cells were controlled by switching between two different gas mixtures: 20% *O*_2_, 5% *CO*_2_ with the balance of *N*_2_, and an oxygen-poor gas mixture: 2% *O*_2_, 5% *CO*_2_ with the balance of *N*_2_, respectively. The flow microchannel, which was 5 mm wide, was connected to a syringe pump (Harvard Apparatus) to control the flow rate (Q) within the microchannel. In this study, the flow rate (Q) ranged from 0 to 20 ml/hr to generate different shear stresses for attaching and detaching the RBCs as required. Imaging of the microfluidic device was carried out using Olympus IX71 inverted microscopes (Olympus, Tokyo, Japan) equipped with Olympus DP72 cameras for image acquisition. All testing was performed at 37°C using a heating incubator (ibidi heating system; ibidi USA, Fitchburg, WI). The microfluidic device was fabricated by permanent covalent bonding of PDMS channels and the glass substrate following air plasma treatment for 1 minute in a plasma cleaner. The PDMS channels were fabricated by casting customized SU-8/Si master molds with a degassed PDMS mixture of base and curing agent (10:1, w/w) using the soft-lithography technique. The macrophages were loaded into the microfluidic device by gently pipetting the cell suspension into the microfluidic channel. The macrophage viability was confirmed by its morphological evidence of no cytotoxicity or apoptosis(for more details see S3 Text).

**Figure 1:**
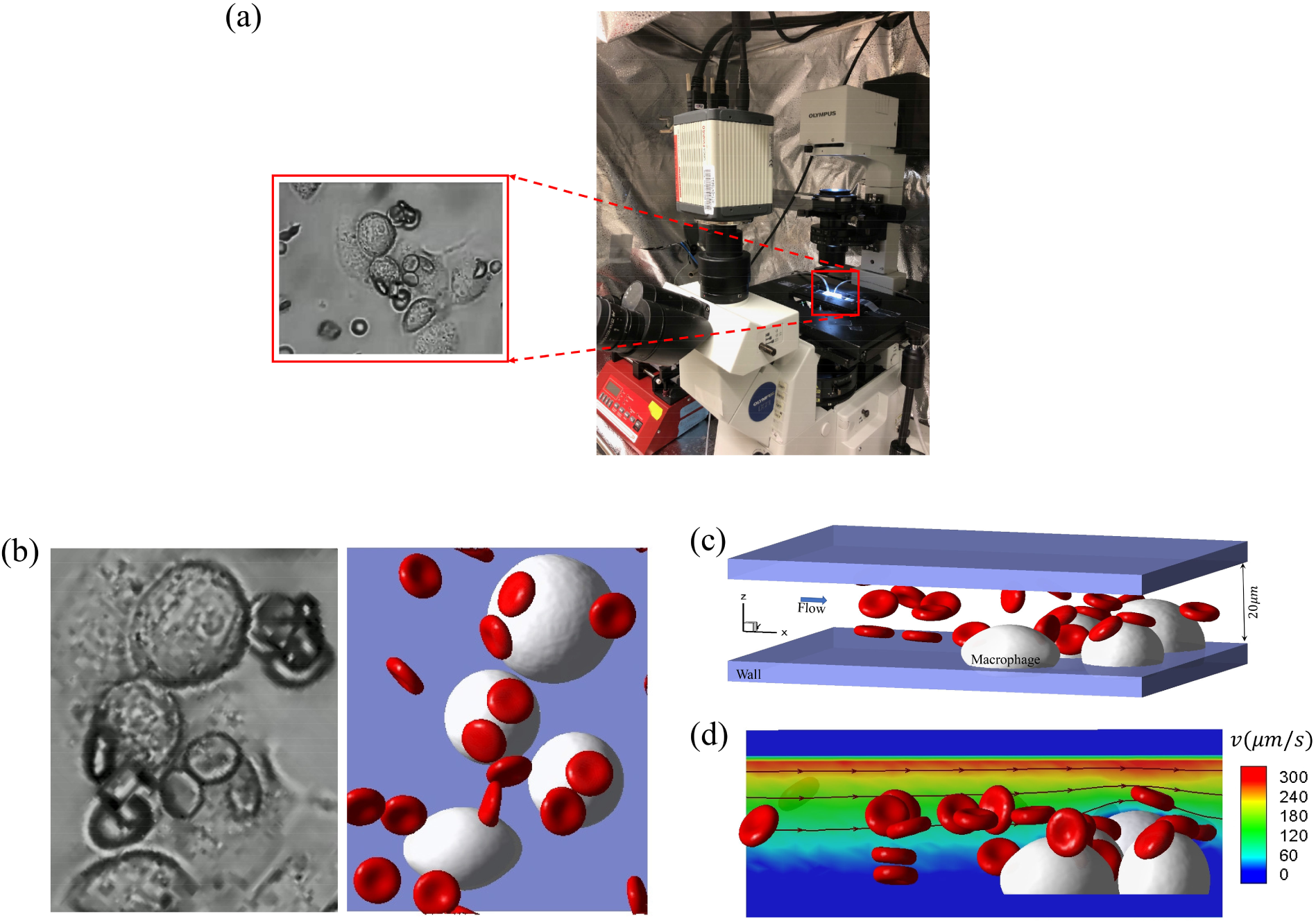
Simulation setup for modeling RBC retention by macrophages under Couette flow. (a) The experiment setup and an observation from the experiment. (b) The snapshots of the cells retained by macrophages in the experiment (left) and simulation (right). (c) The geometry of the simulation setup, where the RBC suspension flows past the macrophages, adhered firmly to the bottom of the chip. (d) Flow streamlines and contours of the velocity field in the simulation. The flow is driven by the velocity of the top wall, while the bottom wall has zero velocity.

It is well-known that macrophages display distinct properties when exposed to hypoxic conditions, such as morphology, migration, chemotaxis, adherence to endothelial cells, bacterial killing, differentiation/polarization, and protumorigenic activity [107–110]. In case of phagocytosis, a prior experimental investigation [111] has shown that hypoxia can enhance the capacity of macrophage to engulf microbeads. Specifically, the phagocytic index is increased by 2, 2.5 and more than 3 folds when the macrophages are treated under hypoxia for 15 mins, 60 mins and 120 mins before phagocytosis experiments, respectively. Since the current study focuses on investigating the impact of hypoxia on sickle RBCs and their adhesion to macrophages, all the adhesion experiments under hypoxic conditions were conducted within a very short time duration (*<*10 minutes), to minimize the impact of hypoxia to macrophages.

#### 2.2.2. Simulation setup

Based on the setup of a typical microfluidic experiment in the left of Fig. 1(b), we construct a companion microchannel *in silico* as shown in the right of Fig. 1(b), where the macrophages are placed on the bottom of a microchannel. The geometry of the microchannel in our simulation is constructed as illustrated in Fig. 1(c), which shows a typical example of simulating the filtration of sickle RBC suspension by a group of macrophages. The height of the microchannel in our simulation is *h* = 20*µm*, and the width and length are determined by the distribution of macrophages and the region of interest recorded in the experiments. We place the macrophages initially on the bottom of the microchannel. Strong adhesion between the macrophages and the channel bottom wall is assumed to ensure no apparent movement of macrophages in our simulation, consistent with the observations from the microfluidic experiments. Since the focus of the simulation is on the adhesion of RBCs to the macrophages, we only simulate the flow around the macrophages (flow field within 20 *µm* from the bottom of the microfluidic channel wall, beyond which these simulated macrophages have no impact on the flowing RBCs or blood flow), instead of the entire channel, to reduce the computational cost. In this case, the height of the simulation domain (20 *µm*) is only 4% of the height of the microfluidic channel. As a result, we approximate the flow field in the simulation domain as a Couette flow. Fig. 1(d) shows the velocity field and streamlines of the flow in the simulation domain. The flow direction in the simulation domain is from left to right, and the top wall’s velocity varies to match the experimental conditions. In this work, we consider the adhesion of sickle RBCs under both normoxia and hypoxia. The shear modulus and bending modulus of sickle RBCs under these conditions are summarized in Table. 2. The shear modulus and bending modulus of normal RBC are chosen to be *E_s_*_0_ = 4.792*µN/m* and *E_b_*_0_ = 2.9 *∗* 10*^−^*^19^*J*, following our prior work in [86]. In addition, as observed from the blood samples of SCD patients, four kinds of shaped cells are considered [30], including biconcave, granular, elongated, and sickle shapes. The normal and sickle RBC models consist of 500 vertices, 996 triangles, and 1494 edges, with a surface area *S_R_* = 132.87*µm*^2^ and a volume *V_R_* = 92.46*µm*^3^. The diameter of macrophages ranges from 16 *∼* 22*µm*, following the measurements from the experiment, and the height of the macrophages is selected to be *h_M_* = 11.00*µm*. Periodic boundary conditions are imposed in the X and Y directions in the simulation, and bounce-back conditions are applied on the channel wall to prevent fluid particles from penetrating.

**Table 2:** Summary of the typical parameters of RBCs under normoxia and hypoxia used in the simulations

| Conditions | Type | $E_s(\text{N/m})$ | $E_b(\text{J})$ | $k_{on}(s^{-1})$ | $k_{off}(s^{-1})$ | $K_s(\mu\text{N/m})$ |
| --- | --- | --- | --- | --- | --- | --- |
| Normoxia | Sickle | $5E_{s0}$ | $5E_{b0}$ | $5.42 * 10^4$ | 2.71 | 2.9 |
| Hypoxia | Sickle | $50E_{s0}$ | $50E_{b0}$ | $8.14 * 10^4$ | 0.54 | 6.7 |

All simulations are computed using the extended version of code developed based on the LAMMPSplatform. Each simulation runs for *∼*3000000 time steps. A typical simulation requires 2304 CPU core hours running on the computational resources (Intel Xeon E5-2670 2.6 GHz 24-core processors) at the Center for Computation and Visualization at Brown University.

## 3. Calibration of the adhesion model based on microfluidic experiments

First, we calibrate the parameters in the adhesion model by matching our simulation results with the experimental observations under normoxia and hypoxia, as the model parameters are not distinguished based on the oxygenation levels in the existing literature. Based on our observations and measurements from microfluidic experiments, we find both normal and sickle RBCs can adhere to macrophages under flow conditions comparable to those in the red pulp of the spleen (*∼* 150 *µm/s*). However, the sickle RBCs under hypoxia detach from macrophages at higher flow velocity than normal and sickle RBCs under normoxia, suggesting a strong adhesive force is triggered for hypoxic sickle cells. The critical detachment velocities, measured at the top of the macrophages, are determined based on the various flow conditions. Typically, the normal RBCs and sickle RBCs under normoxia detach from the macrophages at velocities of approximately 500*µm/s* (video E-3.1, video S-3.1) and 1000*µm/s* (video E-3.2, video S-3.2), respectively. In comparison, a sickle cell under hypoxia has a detaching velocity greater than 2500*µm/s* (video E-3.3, video S-3.3). By matching the simulation results with experimental measurements, we calibrate the key parameters of the stochastic adhesion model, including cell adhesion strength *K_s_*, effective formation strength *σ_on_*, rupture strength *σ_off_*, bond formation rate *k_on_* and bond dissociation rate *k_off_*. We note that the effective formation strength *σ_on_* and rupture strength *σ_off_* have nominal influences on cell adhesion dynamics, so we follow the values used in our previous simulations of sickle RBC adherence to the substrate [98]. Our microfluidic experiments show that the number of RBCs filtrated by the macrophages under hypoxia (video E-3.4, video S-3.4) is approximately five times higher than that under normoxia (video E-3.5, video S-3.5). To match these observations, we tune the parameters *k_on_* and *k_off_*, inspired by our previous study in [98] and find that a higher bond formation rate *k_on_* and lower bond break rate *k_off_*promote the formation of the bonds between RBCs and macrophages. The parameters *k_on_* and *k_off_* are chosen from the range of 5.42*∗*10^4^ *−*8.14*∗*10^4^*s^−^*^1^ and 0.54*−*2.71*s^−^*^1^, respectively, following the values reported in the literature. Eventually, we identify that when *k_on_* = 8.14 *k_off_* = 0.54 under hypoxia state and *k_on_* = 5.42, *k_off_* = 2.71 under normoxia state, our simulation results are comparable with the experimental measurements. More details about the comparison between simulations and experiments will be discussed in Sec. 4.1.2 and 4.1.3. Then, we investigate the adhesion dynamics of normal and sickle RBCs under both normoxia and hypoxia, through which we infer the cell adhesion strength *K_s_* by increasing the flow velocity until the onset of cell detachment. Specifically, we determine the *K_s_* value by fitting the experimental flow velocity values required for cell detachment. A greater *K_s_*will contribute to a stronger cell adhesion strength, resulting in a firm adhesion and a high flow velocity for inducing RBC detachment from macrophages. In comparison, the adhered RBCs can easily detach from the macrophages with a smaller *K_s_*. The details of how we identify the adhesive strength *K_s_* are described in S2 Text. All the calibrated parameters are summarized in Table 2.

## 4. Results

In this section, we integrate computational modeling with microfluidic experiments to quantify the filtrated function of macrophages in microchannels. First, the aggregation of RBCs is introduced to examine the impact of the RBC aggregation force on the macrophage filtration function. Additionally, we investigate the influence of the hematocrit of RBC suspension and blood flow velocity patterns, including constant velocity and step-wise velocities on the adhesion of sickle RBCs to the macrophages. Finally, we present the effects of the morphology of sickle cells on the filtration by macrophages.

### 4.1. Sickle RBC adhesion to macrophages under normoxia state

Fig. 2(a) shows three snapshots of sickle cell suspension adhering to the macrophages under normoxia. The top row displays the experimental observations(video E-3.4), whereas the bottom row depicts the corresponding simulation results(video S-3.4). The configuration of the macrophages in the simulation is derived from the experiments, and the radius range varies from 9 to 11*µm*. The velocity around the macrophages is approximately 150*µm/s*, and the hematocrit is about 2% in the microchannel. Here, *t*_1_ marks the initial stage with no cell flowing through the macrophages, *t*_2_ represents the snapshot when the first RBC adheres to the macrophages, and *t*_3_ is a snapshot when the number of adhered RBCs reaches a plateau. Fig. 2(b) shows the geometry of the microchannel, with the two blue cells representing the adhered cells to macrophages. We also count the number of cells attached to the macrophages over time in the experiment and the corresponding simulation as illustrated in Fig. 2(c), which shows that only two cells are detained by the macrophages from the suspension both in the simulation and experiment within the same period.

**Figure 2:**
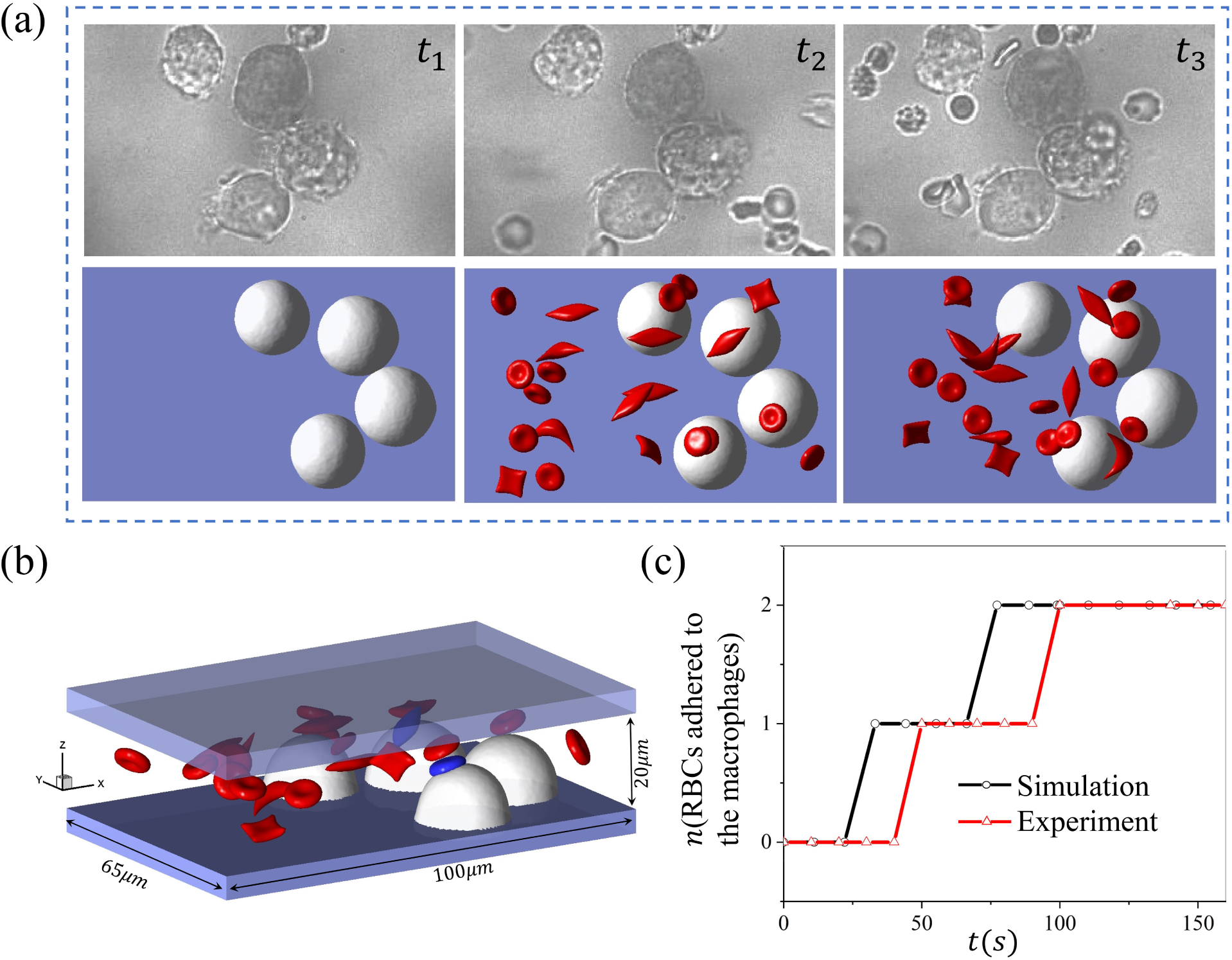
Modeling the sickle RBC retention by macrophages under normoxia. (a) *t*_1_ *∼ t*_3_ shows the three sequential snapshots of RBCs adhering to macrophages from microfluidic experiments (top) and the simulation (bottom). (b) The microfluidic chip’s geometry and the simulation’s final snapshot, with the blue-colored cell adhering to the macrophages. (c) The number of sickle cells retained by the macrophages is recorded over time for microfluidic experiments and simulations.

### 4.2. Sickle RBC adhesion to macrophages under hypoxia state

Fig. 3(a) shows three snapshots of sickle cell suspension filtrated by the macrophages under hypoxia. The experimental observations(video E-3.5) and simulation results(video S-3.5) are illustrated at the top and bottom rows, respectively. The configuration of the macrophages in the simulation is reproduced based on the experimental observations, and the radius varies from 8 to 11*µm*. The velocity around the macrophages is selected to be 150*µm/s*, close to the physiological blood flow velocities in the red pulp of the spleen, and the hematocrit is about 2% in the microchannel. Again, *t*_1_ marks the initial stage with no cell flow through the macrophages, *t*_2_ represents the snapshot when the first RBC adheres to the macrophages, and *t*_3_ is a snapshot when the number of adhered RBCs reaches a plateau. Fig. 3(b) describes the geometry of the microchannel, with the blue-colored cells highlighting the RBCs attached to the macrophages. We also count the number of cells adhered to the macrophages over time in the simulations and experiments as illustrated in

**Figure 3:**
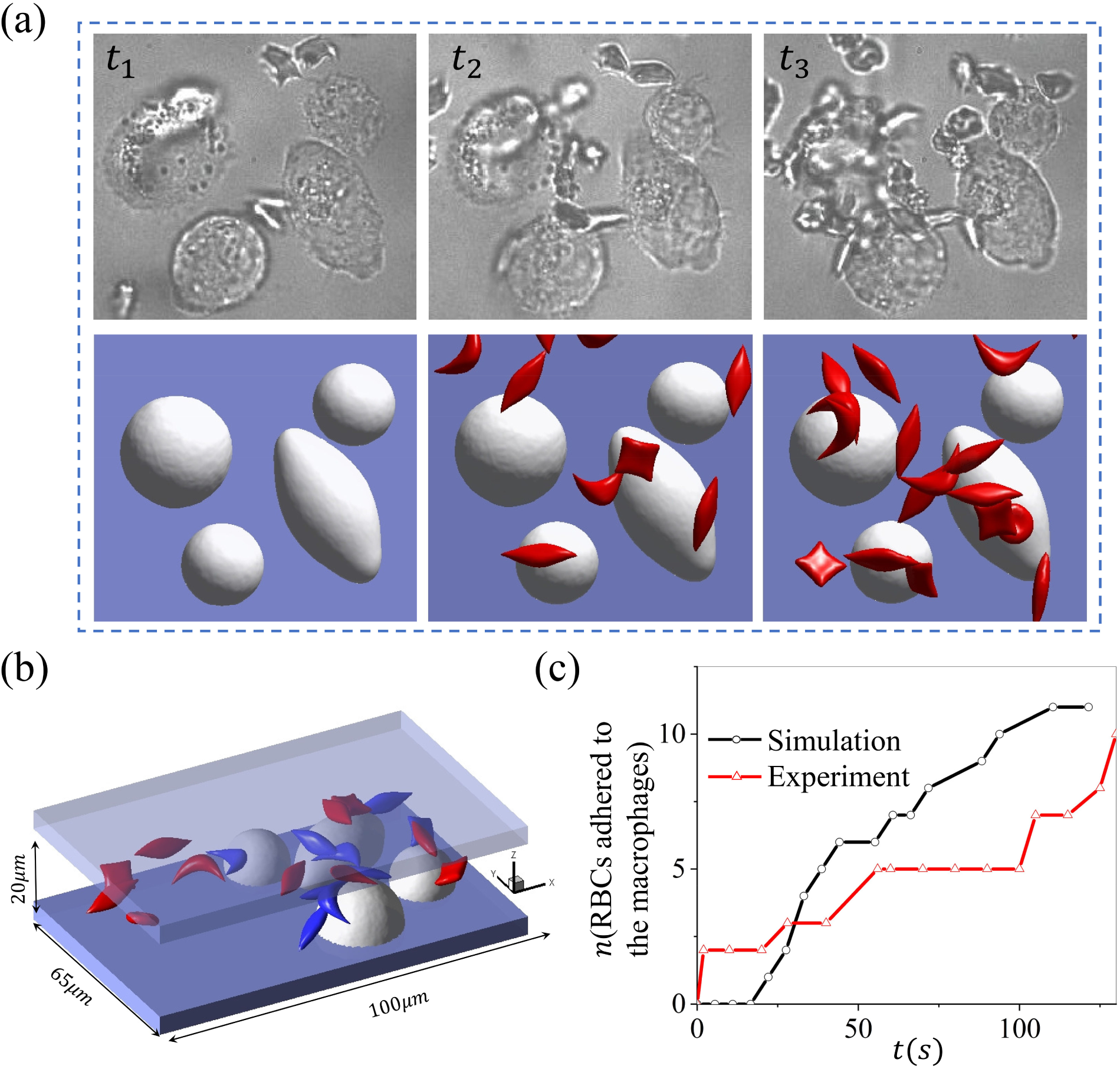
Retention of sickle RBCs by macrophages under hypoxia. (a) *t*_1_ *∼ t*_3_ depicts the snapshots of RBCs adhering to macrophages in microfluidic experiments (top) and simulation (bottom), including the initial (*t*_1_), intermediate (*t*_2_), and final stage (*t*_3_). (b) The microfluidic chip’s geometry and the simulation’s final snapshot, with the blue-colored cells highlighting the RBCs that are firmly adhered to the macrophages. (c) The number of sickle cells retained by macrophages is recorded over time in microfluidic experiments and simulations.

Fig. 3(c), where we note that the number of cells adhered to the macrophages increases over time. The number of cells detained by the macrophages under hypoxia is approximately five times higher than that in the case of normoxia.

### 4.3. Aggregation of RBCs promotes their retention by macrophages

RBC aggregation has been observed in low-shear blood flow where a linear stacking of RBCs can form a chain-like structure, or “rouleau’ [112]. Under pathological conditions such as type 2 diabetes and SCD, the enhanced RBC adhesion due to increased levels of plasma factors like fibrinogen could lead to more pronounced rouleau formation in patient’s blood [113–116], causing blood hyperviscosity in capillaries [117, 118]. Several experimental investigations have been conducted to explore the RBC-RBC aggregation strength using different *in vitro* experimental techniques as well as numerical simulations [119–123]. Despite these aforementioned studies, many important aspects of the influence of cell aggregation on the RBC adhesion to macrophages are still not well understood. Herein, we apply a Morse potential to sickle RBCs in proximity to generate intercellular adhesive force for simulating the aggregation of sickle RBCs. This strategy has been used in many previous works for modeling the formation of RBC aggregates [81, 99, 113, 124]. Fig. 4(a, top row) depicts a process of sickle RBCs detached by macrophages under hypoxia in the microfluidic channel. The first row shows the three snapshots of the experiment(EXP)(video E-4.3). In contrast, the middle and bottom show the simulation results for sickle RBCs with (SIM1)(video S1-4.3) and without (SIM2)(video S2-4.3) considering RBC aggregation. In the simulation, the velocity around the macrophages is about 150*µm/s*, and the radius of macrophages is approximately 11*µm*. Here, *t*_1_ *− t*_3_ depict the snapshots of the entire computational and experimental process, with *t*_1_ and *t*_3_ representing the snapshot of the first cell retained by the macrophages and the final snapshot when the number of adhered RBCs reaches a plateau.

**Figure 4:**
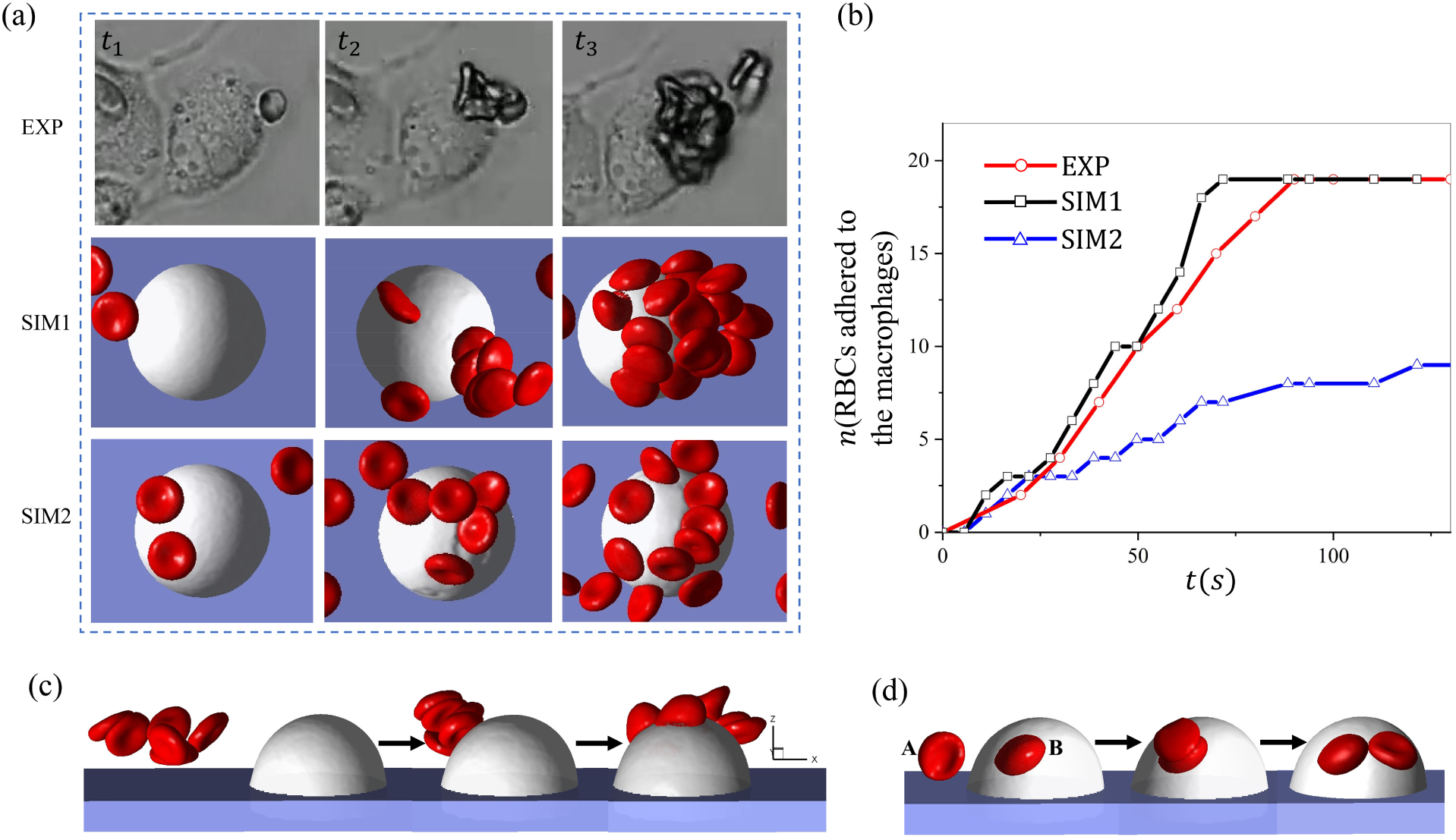
Investigation of the impact of RBC aggregation on RBC retention by macrophages under hypoxia. Snapshot of RBCs adhering to macrophages from experiments and simulations. SIM1 represents the simulations considering the aggregation between sickle RBCs while SIM2 does not. (b) The plot depicts the number of RBCs adhered to the macrophages over time for both experimental and simulation studies. (c-d) The plots describe the two possible adhesion mechanisms that regulate the RBC adhesion to the macrophages (c, cluster filtration) and (d, transitive filtration).

The number of RBCs adhered to the macrophages measured from EXP, SIM1, and SIM2 is plotted in Fig. 4(b), where we observe that the number of RBCs retained by the macrophages computed from SIM1 is comparable to the results of EXP, and twice that of SIM2, indicating the promoting impact of RBCs aggregation to the retention of sickle RBCs by macrophages. Next, we dissect the mechanism of how cell aggregation improves RBC retention. Figs. 4(c) and (d) illustrate two potential mechanisms, which distinguish the filtration results in SIM1 and SIM2, namely cluster filtration (Fig. 4(c)) and transitive filtration (Fig. 4(d)). Cluster filtration occurs when an RBC aggregate approaches and adheres to the macrophage. Multiple RBCs in the aggregate make contact and adhere to the macrophages, leading to a higher retention rate. Under the transitive filtration mechanism, the RBCs adhere to the macrophages one by one; thus, the resulting RBC retention rate is lower than the cluster filtration. As observed from our microfluidic experiments, both mechanisms may occur in the process of RBC filtration by splenic macrophages, depending on the extent of the RBC-RBC adhesion.

### 4.4. Increased Hct of RBC suspension enhances RBC retention by macrophages

In this section, we will examine the effect of Hct of RBC suspension on macrophage retention. We select the Hct to be *Hct* = 1.5%, 2.8%, 3.6%, and 5.0% in our simulations, consistent with the Hct we used in the microfluidic experiments. The variation of Hct is achieved by changing the microchannel length, while the number of RBCs in the simulations remains constant. The radius of the macrophages is 11*µm*, and the velocity is 150*µm/s* around the macrophage. Fig. 5(a) shows the snapshots for these four examined Hct when the number of adhered RBCs in the simulations reaches a plateau. We record the number of RBCs adhered to macrophages over time under different Hct levels in Fig. 5(b), where we observe that although the number of adhered RBCs in all four cases increases with time, simulations with a greater *Hct* takes less time to reach the plateau, as a greater *Hct* could increase the chance of RBCs making contact with macrophages. Additionally, we note that the macrophages retain fewer cells at *Hct* = 1.5%, which is likely because a lower *Hct* diminishes the extent of the RBC interaction that could push RBCs toward macrophages and thus promote RBC adhesion.

**Figure 5:**
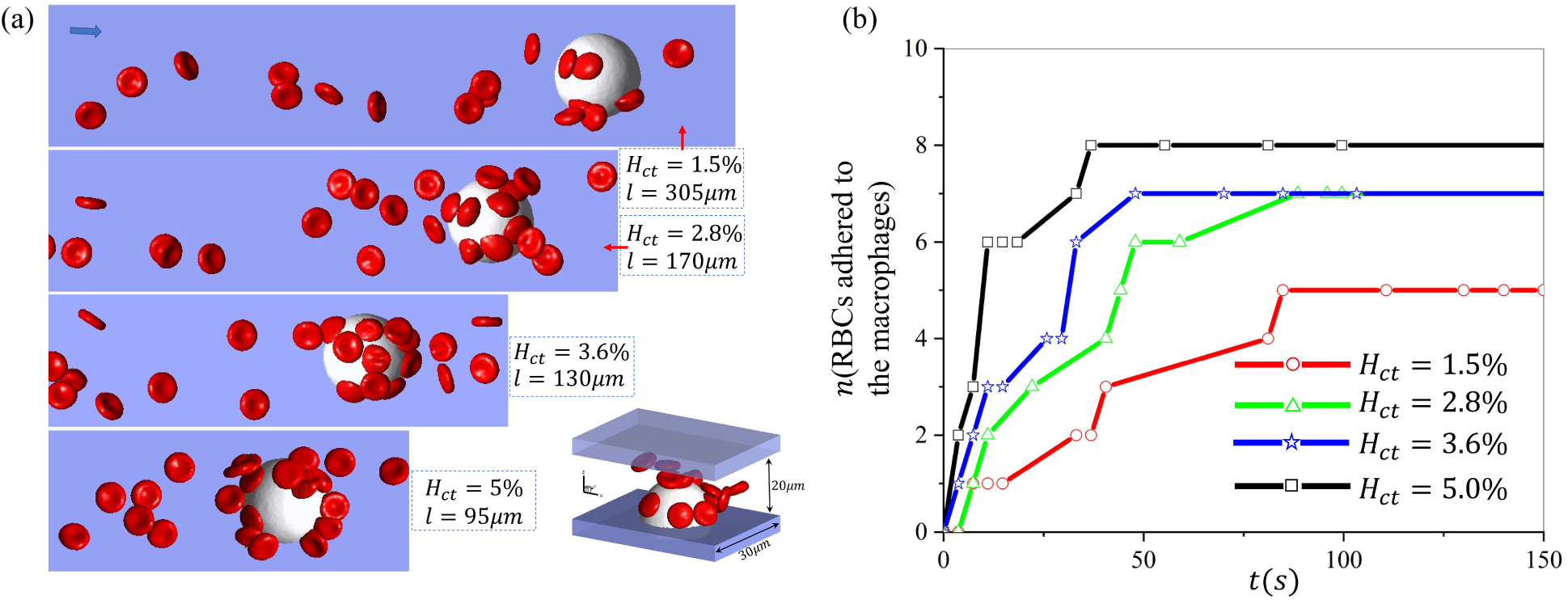
Investigation of the impact of hematocrit on sickle RBC retention by macrophages under hypoxia. (a) Snapshots of RBC suspension with various hematocrits (1.5%, 2.8%, 3.6%, 5%) flowing past a macrophage. (b) The number of RBCs retained by the macrophages is recorded over time under different hematocrits.

### 4.5. Impact of flow velocities on RBC retention by macrophages

Next, we investigate the impact of blood flow velocities on the retention of RBCs by macrophages. Fig. 6(a) depicts the snapshots of RBCs adhering to macrophages at various velocities ranging from 100 *−* 500*µm/s*. The radius of the macrophages in the simulations is 11*µm* and the hematocrit is 1.5%. Fig. 6(b) plots the change of average velocity of the RBCs suspension over time. The results illustrate that the average velocity decreases over time due to the retention of RBCs by the macrophages and eventually reaches a saturated baseline. Fig. 6(c) summarizes the number of cells adhered to the macrophages over time. We note that the number of adhered RBCs rapidly approaches the plateau when a greater velocity is applied in the simulation, which is consistent with the velocity variations in Fig. 6(b). When the velocity is equal to 100*µm/s*, 200*µm/s*, and 300*µm/s*, respectively, the total number of adhered cells is the same when the simulations reach the equilibrium state, while the total number of adhered RBCs at equilibrium decreases as the velocity is increased to 400*µm/s* and 500*µm/s*. These results suggest that the increased blood velocity could attenuate the adhesion of sickle RBCs to the macrophages.

**Figure 6:**
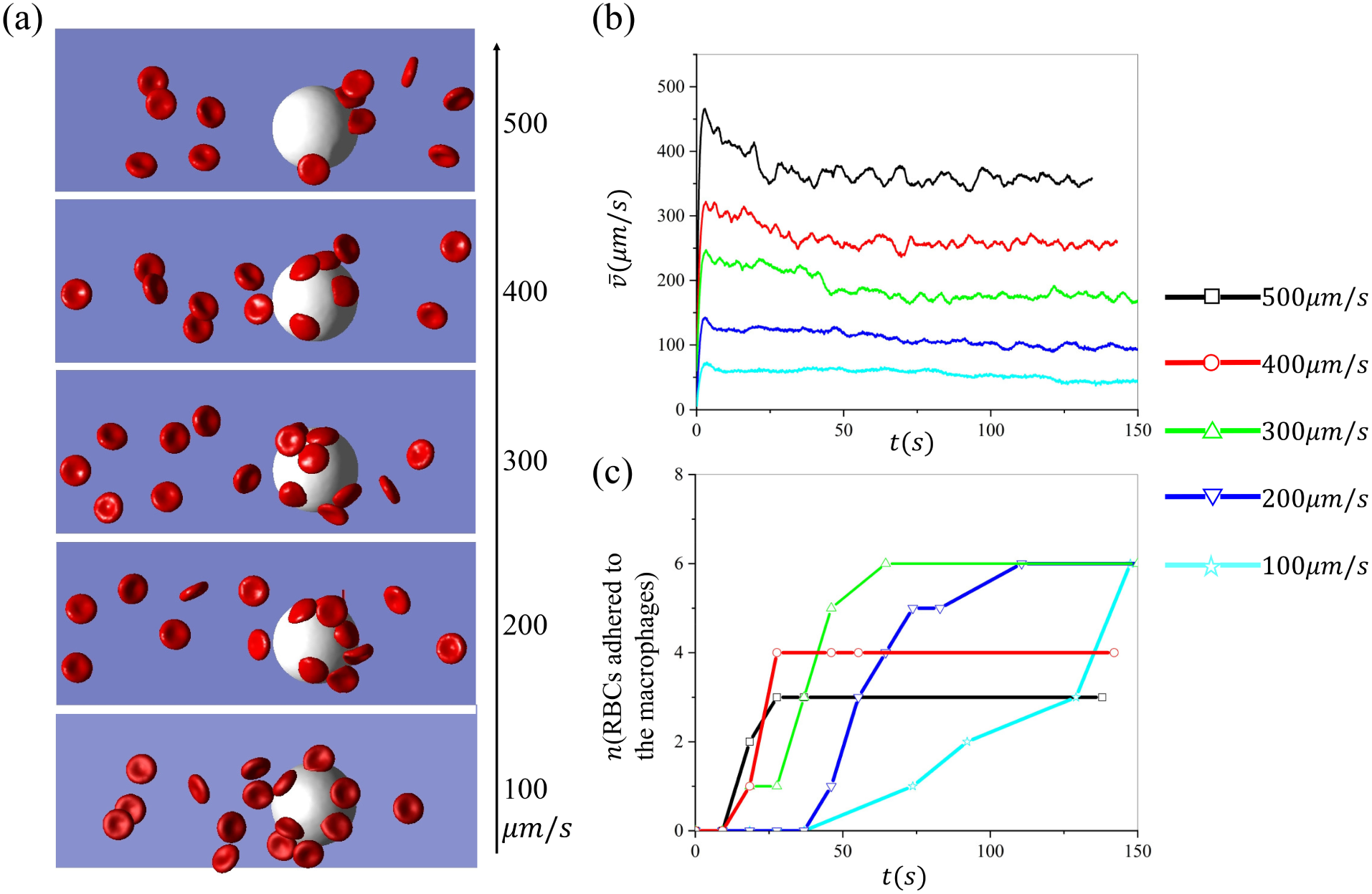
Investigation of the impact of blood velocities on sickle RBC retention by macrophages under hypoxia. (a) The snapshots of sickle RBCs suspension flowing past a macrophage under various velocities. (b) The average velocity of the RBCs suspension changes over time during the retaining process. (c) The number of RBCs retained by the macrophages is recorded over time under various flow velocities.

To further explore the effect of velocity on the RBC retention, we implement a setup that integrates simulation (video S-4.5) and experiment (video E-4.5) to investigate the critical velocity for the RBC adhesion to macrophages using blood velocities regulated in a step-wise manner. The examined velocities are decreased from 1000*µm/s* to 100*µm/s*. Fig. 7(a) shows the consecutive snapshots at different velocities. *t*_1_ is the snapshot when the velocity is *v* = 1000*µm/s* and no RBCs are retained by the macrophage. When the velocity is decreased to about *v* = 500*µm/s*, an RBC is captured by the macrophage at *t*_2_. As the velocity continuously decreases to around *t*_3_(250*µm/s*) and *t*_4_(100*µm/s*), an increased number of RBCs adhere to the macrophage. Fig. 7(b) describes the velocity changes over time in the simulation and experiments, as well as the front view of snapshots of the simulation at various velocities. Following the cell suspension simulations, we perform the single-cell adhesive dynamics study under the velocity *v* = 150, 500, 750*µm/s* in Fig. 7(c) to explain the mechanism causing the difference in the RBC retention at different flow velocities. Our results show that when the velocity is greater than 500*µm/s*, the flowing RBC cannot make contact with the macrophage and they thus cannot be captured and retained by the macrophage. In contrast, a reduced flow velocity will enable contact between the traveling RBCs and the macrophage.

**Figure 7:**
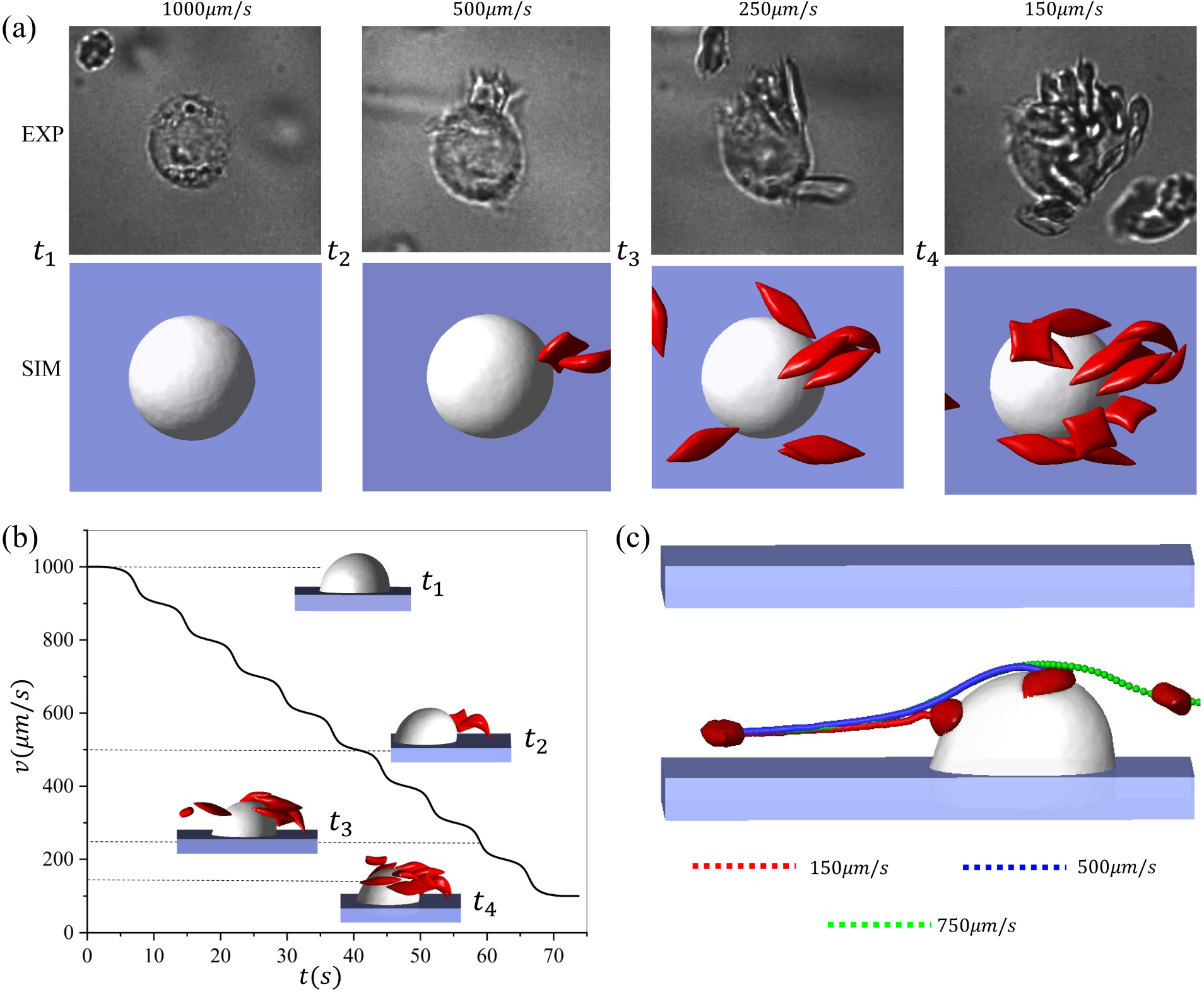
The dynamics of sickle RBCs interacting with a macrophage under a blood flow with step-wise changes. (a) Sequential snapshots of RBCs adhering to macrophages under velocities that are decreased in a step-wise manner. Top: microfluidic experiments. Bottom: the companion simulations. (b) The stepwise velocity changes over time and the front view of the simulation snapshots captured at velocities of *v* = 150, 250, 500, 1000*µm/s*. (c) The impact of the flow velocity on the trajectories of single RBCs flowing past a macrophage.

### 4.6. Impact of the shapes of sickle RBCs on their retention by macrophages

Finally, we investigate the effect of the morphology of the sickle RBCs on their retention by macrophages under hypoxia. Following our prior work in [30], four RBC shapes are considered, including biconcave, elongated, granular, and sickle shapes. Guided by the experimental observation in Fig. 8(a), the radius of the macrophages in the simulation is selected from 8 to 11*µm*. The *Hct* is set to be 2%. Fig. 8(a) shows the snapshots of experiments and simulations with four morphologically altered RBCs. Fig. 8(b) illustrates the changes in the average RBC velocities over time when the four types of RBCs suspension flow past the macrophages at a velocity of 150*µm/s*, where we observe continuous decreases of RBC velocities due to the gradual adhesion of RBCs to the macrophage. Fig. 8(c) records the number of RBCs retained by macrophages for five groups of RBC suspension during the simulation, implying a potential ranking of the likelihood of sickle RBC adhesion to the macrophages: sickle*>*granular*>*mix*>*biconcave *>*elongated. These results suggest that there may be a ranked preference for RBC shapes when macrophages clear sickle RBCs in the spleen.

**Figure 8:**
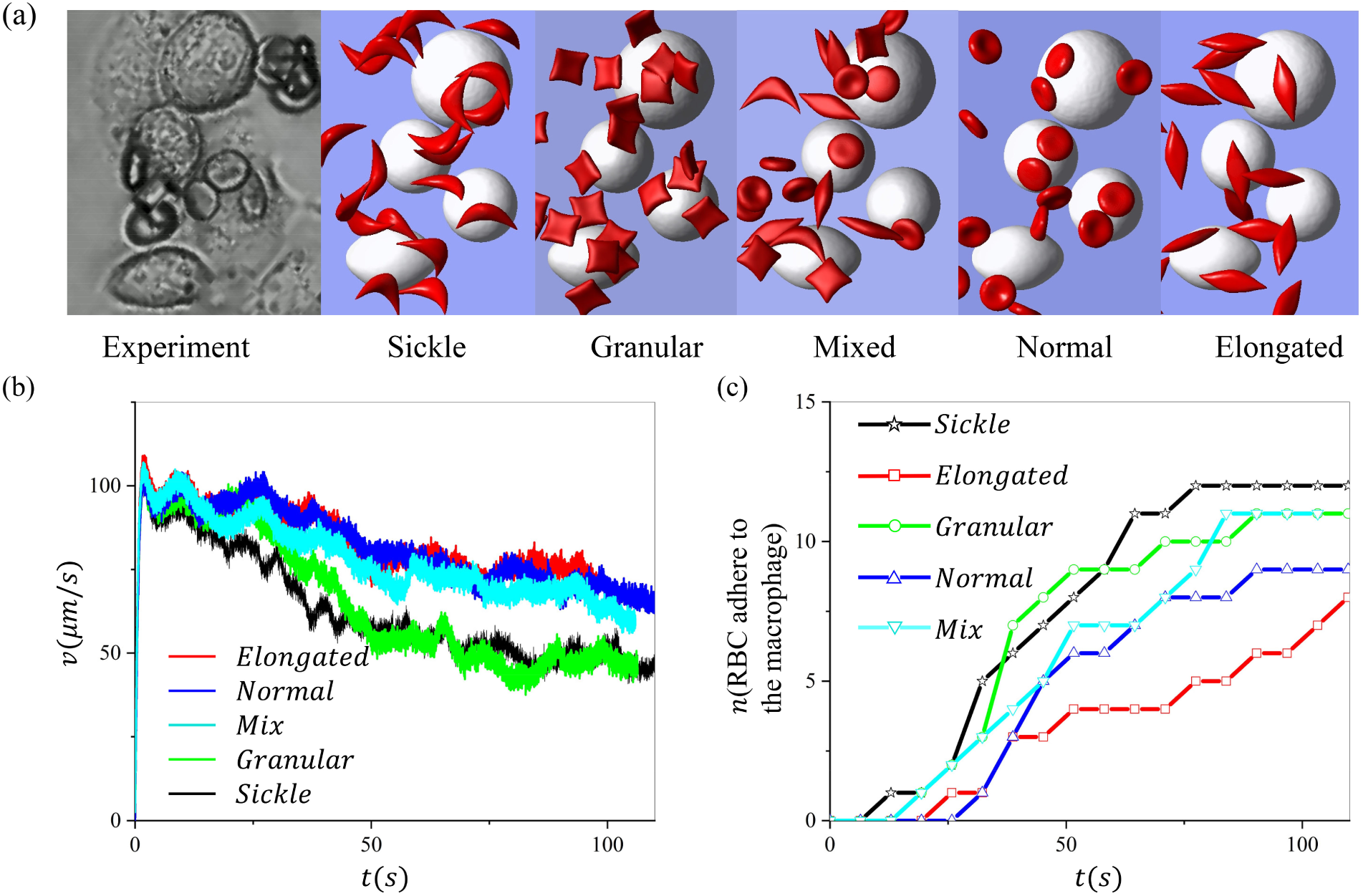
Investigation of the impact of sickle RBC morphology on their retention by macrophages under hypoxia. (a) Snapshots of retention of RBCs with various shapes by macrophages. (b) The average velocity of sickle cells during the retaining process under a bulk flow velocity of *v* = 150*µm/s*. (c) The number of RBCs adhered to the macrophages is recorded over time for sickle RBCs with different shapes.

## 5. Discussion and Summary

In this work, we quantify the retention of sickle RBCs by macrophages by integrating simulations and microfluidic experiments, aiming to improve our quantitative understanding of the filtration function of macrophages in the red pulp of the spleen. Based on the microfluidic experiments, we calibrate the stochastic adhesion model employed to represent the adhesion between the RBCs and macrophages. We focus on identifying the adhesive forces between the sickle RBCs and macrophages under normoxia and hypoxia, which are missing in the literature. By matching our simulation results with experimental observations, we discover that the adhesive strength between sickle RBCs and macrophages is much more robust under hypoxia than under normoxia. As a result, the number of sickle RBCs retained by the macrophages under hypoxia is approximately five times higher than that under normoxia. Notably, this enhanced adhesion strength results in a firm contact of sickle RBCs to macrophages, which can sustain flow velocities more than ten times higher than the flow velocities in the red pulp of the spleen. Although the underlying molecular mechanism causing the increase in the adhesive strength under hypoxia is not yet clear, our results indicate that the filtration function of the splenic macrophages is upregulated under a more hypoxic environment.

Sickle RBCs under hypoxia are featured with high RBC adhesion that facilitates their adhesion to endothelium cells [125, 126], but also to other sickle RBCs [115, 116], both of which could contribute to the initiation and propagation of vaso-occlusion events in SCD. Our microfluidic experiments demonstrate that the formation of RBC aggregate promotes the retention of sickle RBCs by macrophages. The companion simulations we performed further dissect the adhesion process and show that the RBC aggregate adhered to the macrophages could either directly capture the RBCs in the flow or slow down their motion such that the macrophages could retain them. These findings support the role of RBC aggregation in promoting RBC retention by macrophages. Additionally, we examine the impact of Hct on RBC retention by macrophages. Our results show that an increased Hct could boost the chance of RBCs interacting with macrophages, thereby facilitating RBC retention. Considering that the Hct in the red pulp could be as high as 70%, the Hct-induced enhancement of RBC retention could become more pronounced in vivo. We note that although the RBC aggregation and increased Hct promote the retention of RBCs by the macrophages, it is not clear whether the macrophages could efficiently engulf these adhered RBCs. Phagocytosis of RBCs has been primarily studied in vitro and the phagocytic time could vary from minutes to hours [24, 25, 127, 128]. Failure to a timely elimination of these retained RBCs could cause congestion and even ASSC in the red pulp, which has been evidenced by pathological studies of the spleen of SCD patients [43, 129].

We also quantify the impact of blood flow velocities on the retention of RBCs by macrophages by adjusting step-wise the blood flow velocities in the microchannel. Our simulation results show that when the flow velocity is varied between 100-300*µm/s*, which are comparable to the blood velocities in the red pulp, no notable changes occur to the total number of retained RBCs. However, as velocity continues to increase, the number of the retained cells will decrease rapidly, implying that the increased blood velocity could attenuate the function of the red pulp macrophages on detaining aged or diseased RBCs. This finding provides a possible rationale for the slow blood flow in the open circulation of the spleen in facilitating the filtering of RBCs by red pulp macrophages. Our simulations of sickle RBCs with various shapes show shape preference when sickle RBCs are susceptible to retention by macrophages, and the preferential shapes rank in the following order: sickle*>*granular*>*mix*>*biconcave *>*elongated. Our results in the current work suggest that the sickle and granular-shaped RBCs are more likely to be retained in the spleen. This finding is consistent with the observation in the literature that the sickle and granular-shaped RBCs are less frequently observed than the discoid or elongated form of RBCs in the blood smear of SCD patients [30].

We note some limitations both in our experimental results and simulations. In the microfluidic experiments, we used diluted blood samples from SCD patients (Hct*≤*5%) to observe the adhesion dynamics of sickle RBCs to the macrophages in order to quantify the number of adhered RBCs and reduce the risk of clogging the microfluidic channels. This value of Hct used is much smaller than the physiological value of *∼* 70% in the red pulp of the spleen and thus could lead to an underestimation of the function of macrophages on retaining the sickle RBCs. We also did not consider the macrophages’ engulfment of the adhered RBCs, as the phagocytic time is much longer than the simulation time required for forming RBC-macrophage adhesion. We note that modeling the phagocytosis process requires more detailed modeling of intracellular components, such as the polymerization of actin to filaments and the interaction between the actin filaments and macrophage membrane, which is beyond the capability of the models employed in the current study. We use a general kinetic cell adhesion model proposed by Bell and Dembo[130, 131] to mimic the collective effect of ligand-receptor binding between the sickle RBCs and macrophages. This model cannot distinguish the contribution from various ligands, such as the band-3 proteins [17, 18], exposure of PS [19, 20], decreased expression of CD47 [21, 22] and conformational changes in CD47 [21, 23].

In summary, we have employed mesoscopic computational models that are validated by microfluidic experiments to simulate the adhesion of sickle RBCs to macrophages, aiming to infer the filtration function of the red pulp macrophages. This important spleen function has not been well-studied. We quantified the impact of several key factors that could dictate RBC retention by splenic macrophages, such as blood flow conditions, RBC aggregation, Hct, RBC morphology, and particularly oxygen levels, which, to the best of our knowledge, have not been systematically explored before. Our experimental and simulation results improve our quantitative understanding of the filtration function of splenic macrophages and provide an opportunity to combine the new findings with the current knowledge of the filtration function of IES to apprehend the complete filtration function of the spleen in SCD.

## Author Contributions

G. L.: designed research, performed research, contributed analytic tools, analyzed data and wrote the paper.

Y. Q.: performed research, analyzed data and wrote the paper.

H. L.: designed research, contributed analytic tools, analyzed data and wrote the paper.

X. L.: contributed analytic tools, analyzed data and wrote the paper.

P. A. B.: designed research, analyzed data and wrote the paper.

M. D.: designed research, analyzed data and wrote the paper.

G. E. K.: designed research, analyzed data and wrote the paper.

## Declaration of Interest

The authors declare no competing interests.

## Acknowledgment

This work was supported by the National Heart, Lung, and Blood Institute of the National Institute of Health under grant number R01HL154150. High Performance Computing resources were provided by the Center for Computation and Visualization at Brown University.

## Notes

### Competing Interest Statement

The authors have declared no competing interest.

